# ExpertRNA: A new framework for RNA structure prediction

**DOI:** 10.1101/2021.01.18.427087

**Authors:** Menghan Liu, Giulia Pedrielli, Erik Poppleton, Petr Šulc, Dimitri P. Bertsekas

## Abstract

Ribonucleic acid (RNA) is a fundamental biological molecule that is essential to all living organisms, performing a versatile array of cellular tasks. The function of many RNA molecules is strongly related to the structure it adopts. As a result, great effort is being dedicated to the design of efficient algorithms that solve the “folding problem”: given a sequence of nucleotides, return a probable list of base pairs, referred to as the secondary structure prediction. Early algorithms have largely relied on finding the structure with minimum free energy. However, the predictions rely on effective simplified free energy models that may not correctly identify the correct structure as the one with the lowest free energy. In light of this, new, data-driven approaches that not only consider free energy, but also use machine learning techniques to learn motifs have also been investigated, and have recently been shown to outperform free energy based algorithms on several experimental data sets.

In this work, we introduce the new ExpertRNA algorithm that provides a modular framework which can easily incorporate an arbitrary number of rewards (free energy or non-parametric/data driven) and secondary structure prediction algorithms. We argue that this capability of ExpertRNA has the potential to balance out different strengths and weaknesses of state-of-the-art folding tools. We test the ExpertRNA on several RNA sequence-structure data sets, and we compare the performance of ExpertRNA against a state-of-the-art folding algorithm. We find that ExpertRNA produces, on average, more accurate predictions than the structure prediction algorithm used, thus validating the promise of the approach.

## 1 Introduction

Ribonucleic acid (RNA) is a fundamental biological macromolecule that is essential to all living organisms, performing a versatile array of cellular tasks including information transfer, enzymatic function, sensing, regulation, and structural function (Elliott and Ladomery, 2017). RNA has recently emerged as a promising drug target, with new therapeutic approaches aiming to develop drugs that target RNA rather than proteins. Moreover, designed RNA molecules are used in rapidly growing fields of synthetic biology and RNA nanotechnology, with applications to diagnostics, immunotherapy, drug delivery and realization of logical operations inside cells (Green et al., 2014; Geary et al., 2014; Hochrein et al., 2013; Han et al., 2017; Guo, 2010; Qi et al., 2020). In particular, such functionalities are enabled by the 3D structure the RNA molecules fold into. Each RNA molecule is made up of individual units, nucleotides (bases), that are of four types, A, U, G and C. A molecule is then uniquely defined by its sequence and structure. The sequence is a string with the A, U, C, G bases reported in the order they appear. RNA sequences range in length from tens (tRNAs, siRNAs) to tens of thousands (e.g. viral genomes and long non-coding RNAs) of nucleotides. The structure, i.e., the way nucleotides are interacting in space, can be represented through an incidence matrix (each cell is (1,0) if the nucleotides are linked or not, respectively) or dot-bracket notation (Sect. 3). In particular, the primary structural elements of RNA molecules are base pairs, i.e., bonds between bases that are not contiguous. The most common types of base pairings, referred to as the “canonical base pairs”, are (i) the Watson-Crick base pairs, where a nucleotide of type A pairs with U, G pairs with C, and (ii) the wobble base pair, where a nucleotide of type G pairs with U. Given a sequence, a set of canonical base pairs defines the secondary structure, which is often used as a first approximation of the RNA structure. The tertiary structure is a more accurate representation that also considers weaker interactions compared to the base pairing. Together, these interactions determine the three-dimensional relative orientation of nucleotides.

Given the impact of structure on RNA functionality, the accurate computational prediction of the secondary and tertiary structure of RNA is an ongoing area of great interest in the computational biology community (Calonaci et al., 2020; Cruz et al., 2012; Wayment-Steele et al., 2020).

### Secondary Structure Prediction

Most tools for secondary structure prediction (Reuter and Mathews, 2010; Hofacker, 2003; Zadeh et al., 2011; Zuker and Stiegler, 1981) attempt to identify the structure that minimizes the free energy (FE) associated with the RNA molecule upon pairing a subset of the nucleotides, i.e., the energy released by folding a completely unfolded RNA sequence. The underlying assumption is that the structure with the lowest free energy is also the most likely structure the RNA will adopt. Equivalently, this family of approaches relies on the basic idea that the lower the FE the more stable the RNA structure will be. A *first* challenge for this family of approaches is that it is not possible to exactly calculate the free energy due to the (i) incomplete understanding of the RNA molecular interactions, and (ii) the impractical computational cost of detailed kinetic simulation tools. As a result, several approximate models have been proposed in the literature (Mathews et al., 1999; Xia et al., 1998; Andronescu et al., 2007, 2010) to estimate the free energy associated with a given secondary structure. Most of the computational savings are a result of ignoring tertiary interactions. A *second*, and possibly deeper, challenge is that this model assumes that an “optimal” structure is one that pairs nucleotides in a way that minimizes the free energy (MFE). However, RNA is known to fold cotranscriptionally (Angela et al., 2018), equivalently, RNA molecules might adopt a kinetically-preferred structure different from the global free energy minima.

In light of these challenges, alternative approaches to structure prediction have been proposed. Stochastic kinetic folding algorithms (Isambert and Siggia, 2000; Sun et al., 2018) approximate the folding kinetics of RNA molecules as they are transcribed. Data driven approaches have also started to become popular that use machine learning to evaluate structures rather than FE or kinetic models. These include ContraFold, DMfold, and structure prediction with neural networks (Do et al., 2006; Wang et al., 2019; Calonaci et al., 2020). Furthermore, in the attempt to achieve advantages of model or data driven approaches, methods have been proposed that attempt to aggregate multiple information sources to get more accurate secondary structure prediction. Within the data driven category, some examples of information sources are the experimentally determined SHAPE data (Lucks et al., 2011; Low and Weeks, 2010), and evolutionary covariation information (Calonaci et al., 2020). On the model driven side, statistical ensemble approaches are used to boost the solutions obtained by the different FE driven folding algorithms. To the knowledge of the authors, ensemble methods allow to mix solutions from different algorithms only upon completion, i.e., they do not enable interaction among the algorithms while they are running (Aghaeepour and Hoos, 2013). A recent survey on a set of secondary structure prediction tools has reported mixed results, with data-driven approaches generally outperforming the ones based on nearest-neighbor free energy models, and with model-based ensemble approaches showing competitive results (Wayment-Steele et al., 2020).

### Tertiary Structure Prediction

Concerning the tertiary structure prediction problem, fewer approaches can be found in the literature (Seetin and Mathews, 2011; Westhof and Auffinger, 2006; Tan and Chen, 2010; Miao et al., 2017; Watkins et al., 2020). In fact, the prediction of tertiary structures is particularly challenging, and most prediction methods only work for short RNA sequences (tens of nucleotides). Data driven approaches have attracted attention also for the tertiary structure prediction. However, their accuracy remains limited due to the small number of 3D RNA structure data sets available for model training and verification.

Stemming from the observation that several folding algorithms have been proposed in the literature for secondary and, even if fewer, for tertiary structure prediction, without any approach dominating the other, we propose the idea to build a framework that can exploit several folding tools and criteria to evaluate the quality of a folded sequence, *during* the algorithm execution. The aim of our approach is to achieve a better RNA structure prediction quality. The result of our work is ExpertRNA, which allows to combine multiple secondary structure prediction algorithms (equivalently referred to as folding algorithms or folders), whose predictions are sequentially evaluated by a set of *experts* that score the quality of the predicted structures based upon their own scoring criteria. ExpertRNA is based on rollout (Bertsekas, 2020), an approximate dynamic programming technique, which, given an algorithm called base policy (or base heuristic), generates an improved algorithm, called rollout policy, as evaluated by one or more “experts”. Rollout also allows the use of multiple base policies and it provably improves (relative to the expert score) on the performance of each of them.

We argue that ExpertRNA has the potential to exploit the strengths of the folding tool being used by the algorithm itself as base heuristic, by improving on the solution by using several experts. In this manuscript, we focus on secondary structure prediction, and we use RNAfold (a free energy based folder (Lorenz et al., 2011)) as the base heuristic (Sect. 2), and, as expert, we use two different implementations of the state-of-the-art tool ENTRNA (a classifier that evaluates the quality of a sequence-structure pair (Su et al., 2019)) to judge sequence-structure pairs generated by a folding algorithm (Sect. 2). We test ExpertRNA against two popular RNA sequence-structure data sets: Rfam (Burge et al., 2013), and the Mathews data set (Ward et al., 2017). We compare the performance of ExpertRNA against RNAfold, when used in isolation. We find that, in the sequences taken from Rfam, ExpertRNA produces, on average, more accurate predictions of secondary structure, and performs, on average, the same on the Mathews data set.

## 2 Relevant Literature

With respect to our work, we distinguish two main branches of research within the rich literature in RNA secondary structure prediction: (i) *model driven*, and (ii) *data driven* approaches. As mentioned in Sect. 1, most of model driven approaches use the Free Energy (FE) as reference reward function to be minimized. Algorithms differ in the model used to approximate the FE and the mechanisms to explore the space of possible structures, in the attempt to find the minimum FE (MFE). Data driven approaches use machine learning techniques to evaluate the quality of the structure as a replacement, or, possibly, in addition to FE. In this case, FE is interpreted as *a* feature instead of *the* reward.

### 2.1 Model driven folding

The most common approach for evaluating RNA structures is its free energy, which can be thought of as the energy released by folding a completely unfolded RNA sequence. It can also be interpreted as the amount of energy that must be added in order to unfold a folded molecule. As a result, a sequence will be most likely found in a minimum free energy structure. No closed form is available for the exact free energy calculation, and algorithms differ in the way they approximate such computation, and the way they search in the space of secondary structures attempting to find the one(s) with associated minimum (approximated) free energy (Lorenz et al., 2011). An example of minimum free energy approximation relies on the nearest-neighbor model (NN-M) (Xia et al., 1998). The NN-M relies on the assumption that each base pair contributes to the overall free energy independently from one another, and a base pair is only influenced by the immediately adjacent base pairs. As a result of these assumptions, the total free energy is the sum of all the base pair free energies, along with additional terms that account for the free energy contribution of motifs such as hairpin loops, internal loops, and bulges and junctions (Mathews et al., 1999) (Fig. 1). Using the free energy minimization criteria, any predicted “optimal” secondary structure for an RNA molecule depends on the model of folding and the specific folding energies used to calculate that structure. A consequence of this is that different optimal structures may be obtained if the folding energies are changed even slightly.

**Figure 1:**
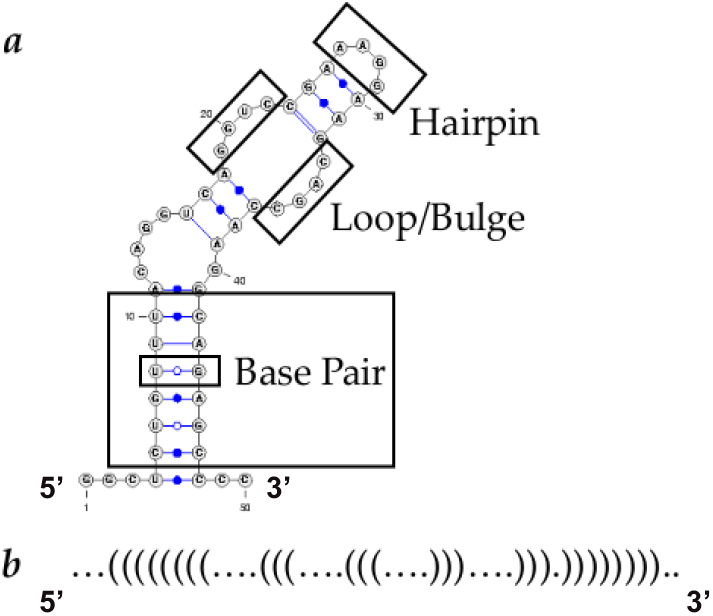
Example of an RNA secondary structure representation, with the highlighted structural motifs: a) 2D planar structure representation; b) Dot-bracket notation. This notation only represents the base pairing and does not provide the type of nucleotide.

An example of algorithm relying on FE is RNAfold, a de-facto standard tool for RNA secondary structure prediction. The RNAfold software (Lorenz et al., 2011) makes use of the NN-M to sequentially assign nucleotides to the partially folded sequence. Importantly, RNAfold allows *constrained folding*, i.e., users can input information concerning pairings between nucleotides that they wish to fix and/or just constraining nucleotides to be paired without defining the nucleotide to pair it with. This second type of constraint is not generally implementable by folding packages and led us to choose RNAfold as the base heuristic for this first version of ExpertRNA. Further examples of model based RNA folding approaches are stochastic kinetic folding algorithms. Rather than using dynamic programming, base pairings are randomly sampled biasing towards bonds with high energy. Energy is estimated using kinetic models, and, for computational efficiency, the kinetic model is called only to evaluate the partially folded structure (Isambert and Siggia, 2000; Sun et al., 2018; Zadeh et al., 2011). Another approach is AveRNA, an ensemble-based prediction method for RNA secondary structure (Aghaeepour and Hoos, 2013). AveRNA combines a set of existing MFE-based secondary structure prediction procedures into an ensemble-based method aiming to achieve higher prediction accuracy. The underlying algorithms use different models for free energy. These models, first presented in (Andronescu et al., 2007, 2010), use the constraint generation method to sequentially produce constraints that enforce known structures to have energies lower than other structures for the same molecule. Such method has resulted in several FE approximation variants.

It is important to, again, highlight that one of the main drawbacks of model driven methods is that energy models are approximations of the not fully known complex physical interactions and folding conditions that determine the folded structure of the molecule in natural or laboratory settings. Because of uncertainties in the folding model and the folding energies, the “true” folding may not be the “optimal” folding determined by the algorithm. In fact, several and sub-optimal structures may exist within a few percent of the minimum energy.

### 2.2 Data-driven folding

Approaches utilizing statistical learning have recently received increasing attention. Different folding algorithms may use different statistical methods and different features to generate base pairing decisions. The CONTRAfold algorithm falls into this category of approaches (Do et al., 2006). CONTRAfold applies stochastic context-free grammars (SCFGs) for representing RNA structures as random objects that are modified within a fully-automated statistical learning procedure that sequentially biases the sampling distribution responsible for the making and breaking of base pairs within the sequentially formed RNA structure. In particular, CONTRAfold sequentially forms and brakes base pairs updating the probability associated to the several bonding alternatives, which is formulated using conditional log-linear models (CLLMs). The hyperparameters of the CLLMs are estimated using several features of RNA sequence-structure pairs. The model determines the likelihood of a base pairing (or a set of pairings) to be selected by the algorithm (Do et al., 2006). The algorithm CycleFold (Sloma and Mathews, 2017) differs from ContraFold because it uses sets of RNA bases as building blocks for the search, instead of the individual nucleotides. Each of these sets has a detailed energetic characterization, thus improving the accuracy of the energy prediction patterns. The building blocks are subsequently folded in a way similar to other data-driven approaches (Dieckmann et al., 1996). LearnToFold (Chowdhury et al., 2019) uses an approximate folding tool (LinearFold) as inference engine for search, and a structured support vector machine (sSVM) to bias the sampling towards more favorable pairings. SPOT-RNA (Singh et al., 2019) uses deep contextual learning for base pair prediction and it is one of the few to include tertiary interactions, even if approximated. Deep and transfer learning (Hanson et al., 2020) are trained in order to calculate the likelihood of each nucleotide to be paired with any other nucleotide in a sequence. ENTRNA (Su et al., 2019) is a supervised learning algorithm that uses a support vector machine (SVM) to produce an evaluation of the quality of a RNA secondary structure. In particular, ENTRNA receives as input a sequence-structure pair and it returns a measure of the likelihood that such a pair folds, and it is stable. Such likelihood measure is defined by the authors as *foldability*. The higher the foldability, the more likely the sequence-structure pair is to exist and be stable. Alternatively, the more likely that sequence is to fold into that structure. ENTRNA, as any data-driven approach, needs to be trained by examples of existing sequence-structure pairs, each associated with a set of features (sequence-segment entropy −SSE-, free energy, relative number of nucleotides of type G and C, relative number of paired nucleotides). In particular, the SSE metric was proposed in (Su et al., 2019) to give a measure of diversity of the nucleotides segments within the RNA sequence. However, to be trained, ENTRNA *also* requires failed examples (i.e., sequence-structure pairs that cannot fold in a stable manner). This is a challenge due to the unavailability of “failed” experiments, which the authors tackle by using the Positive-Unlabeled learning method to fill in the failed examples (Li et al., 2009). Nevertheless, this remains a source of inaccuracy for the approach. Similar to ENTRNA, RNAStructure (Reuter and Mathews, 2010) uses both data driven features and free energy models to evaluate sequence-structure pairs. RNAstructure, first, contained a method to predict the lowest free energy structure, it was subsequently expanded to include bimolecular folding and hybridization thermodynamics, and stochastic sampling of structures. It provides methods for constraining structures based on empirical characteristics of similar sequences. Finally, recent extensions calculate partition functions for secondary structures common to two sequences and can perform stochastic sampling of common structures.

### 2.3 Contributions

The contribution of this paper is both algorithmic and applied: (i) we extend the rollout approach to be used with multiple heuristics and multiple experts, while controlling the number of branches being active at each iteration of the algorithm, thus allowing to control the computational expense. We allow the different experts to interact which leads to better solutions than allowing experts to only judge the branches dedicated to them (Bertsekas, 2020); (ii) we exploit this algorithmic capability to maintain multiple parallel rollout branches allowing the several experts and, potentially, folding algorithms, to maintain multiple solutions. To our knowledge, unlike all the previously published algorithms, ExpertRNA is the only approach allowing the different folding algorithms to interact *while* the structure is being folded. We believe that this capability is the key to overcome the different challenges from the several secondary structure prediction approaches (model and data-driven).

## 3 Proposed Approach

The main idea underlying ExpertRNA is to formulate secondary structure prediction (folding) as a sequential assignment problem. Based on the description in Sect. 2, we assume a sequence of nucleotides is given as input together with a set of physical constraints that define possible nucleotide-to-nucleotide interactions (base pairing) that can be translated into feasible actions. It is important to formalize the way we model RNA sequences and the related structures before introducing actions and constraints.

Specifically, our work makes use of the dot-bracket notation to represent RNA structures (Antczak et al., 2018). According to this formalism, each nucleotide is represented by a letter, symbolizing the type of base (A, C, G, U) and a “dot” (.), if the nucleotide is unpaired, i.e., it does not establish a bond with any other nucleotide in the sequence, or a “bracket” otherwise. If a nucleotide pairs with another, it can either be the origin of the pairing, in which case we represent it using an open bracket “(“, or the sink, in which case we use a close bracket “)” (Fig. 1). Fig. 1 shows an example of an RNA sequence-structure (Fig. 1a), and the corresponding dot bracket notation (Fig. 1b). In the following, we define the main components of ExpertRNA.

### 3.1 State and action modeling in ExpertRNA

As previously mentioned, ExpertRNA sequentially attaches bases to a partially formed structure deciding whether to pair a base (open or closed bracket) or not. Given the same sequence, a different dot-bracket assignment will result in a different structure.

More specifically, when we select the next element from the input sequence *x* ∈ {*A, U, C, G*}, we will connect it to the rest of the structure by means of 3 alternative actions (represented as the top, bottom, and right arcs from each node in Fig. 2). The right arc, used in isolation, corresponds to a dot in the dot bracket notation, and establishes a sequential link between bases (i.e., no base pair), while the up/down (open/closed) arcs establish a base pair type of connection, thus corresponding to a bracket. The number of links that can be activated at iteration *k* is a function of the partial structure formed by the *k* nucleotides already assigned, as well as the remaining *N – k* nucleotides in the sequence that have not been assigned yet. Fig. 2 reports an illustrative example of the sequential folding of the RNA molecule.

**Figure 2:**
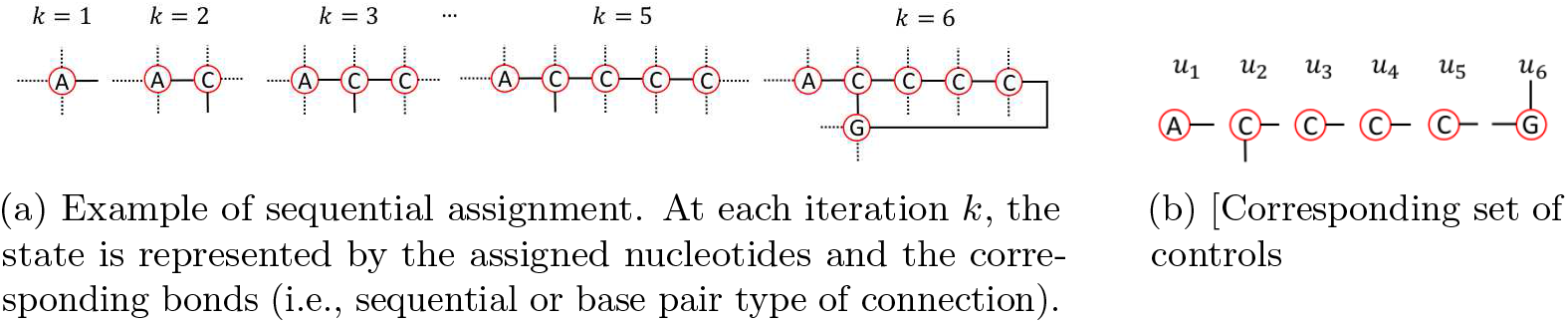
Example of assignment

We start selecting a nucleotide (A in the example in Fig. 2a) and we activate the right link, corresponding to a At the second iteration, we select C, the next nucleotide in the sequence, and we activate the downward link (open bracket “(“)), and the right link, thus electing C as part of a base pair. Note that, at this point, the sink of the base pair is not established, but we have only decided that C will be part of a base pair, as origin. This means that, further in the path generation, we will need to elect one of the candidates to be paired with C. The third-to-fifth actions are identical and they result in three C-type bases with the right link activated, corresponding to three dots (“.”). At this point, we select G as the next base with the upward link active (closed bracket “(”)), thus electing G as base pair, and we connect it to the, only feasible origin C.

#### Actions

Generalizing the example in Fig. 2, the possible actions that we can implement, at each iteration, are to sequence, open base pair and close base pair, with the associated dot-bracket notation respectively. ExpertRNA proceeds to form a structure starting from from the unpaired 5’ extreme (the “first” nucleotide in the sequence, Fig. 1), and completed once all the elements are assigned, i.e., the 3’ extreme of the sequence has been reached (the “last” nucleotide in the sequence, Fig. 1). Also, we assume that bases assigned and links activated at iteration *h* cannot be changed at any iteration *k > h*. Let us consider a sequence of actions {*u*_1_,…, *u_k_*} corresponding to the assignment of the first *k* nucleotides out of a *N*-nucleotides long sequence, with *k* < *N*. Equivalently, {*u*_1_,…, *u_k_*} is a partial assignment. Our objective is to sequentially perform sequence-structure assignments using a set of well defined elementary actions. A structure is complete when *k* = *N*.

#### Feasibility Determination

As previously mentioned, at each iteration of the algorithm, we can either execute a (i) “null” action (sequencing), or (ii) base pairing the nucleotide. When evaluating the feasibility of an action the following items need to be considered:

- nucleotides of type A can only form a base pair with U;
- nucleotides of type G can only form a base pair with C or U;
- nucleotides of type U only form a base pair with A or G;
- nucleotides of type C only form a base pair with G;
- to be paired *i* and *j* need to satisfy |*i* – *j*| ≥ 4, where *i* and *j* are the location indexes of the two nucleotides.

Hence, at each iteration, the set of feasible actions depends on the partial sequence 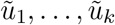 generated by the procedure up to iteration *k*, as well as the nucleotides *k* + 1,…, *N* that remain to be sequenced.

### 3.2 ExpertRNA Algorithm

The proposed ExpertRNA tackles the problem of RNA secondary structure prediction as a sequential assignment of (known) nucleotides.

Fig. 3 shows the structure of our ExpertRNA approach with its two main algorithmic components: (i) the partial folding; and the (ii) expert software. The algorithm sequentially adds elements to the incomplete structure (“current partial folding” in Fig. 3), which we initialize to be the empty set. The first nucleotide is chosen as the first element of the input sequence provided by the user. At each step, the subsequent nucleotide is selected, and we can choose whether to simply *sequence* it to the last assigned nucleotide (“Null” action in Fig. 3) or pair it with any nucleotide in the existing structure (“Close” in Fig. 3), or pair it with an element still to be assigned (“Open” in Fig. 3). The definition of these actions is motivated by the physical laws that govern molecular bonding (as previously specified in feasibility determination).

**Figure 3:**
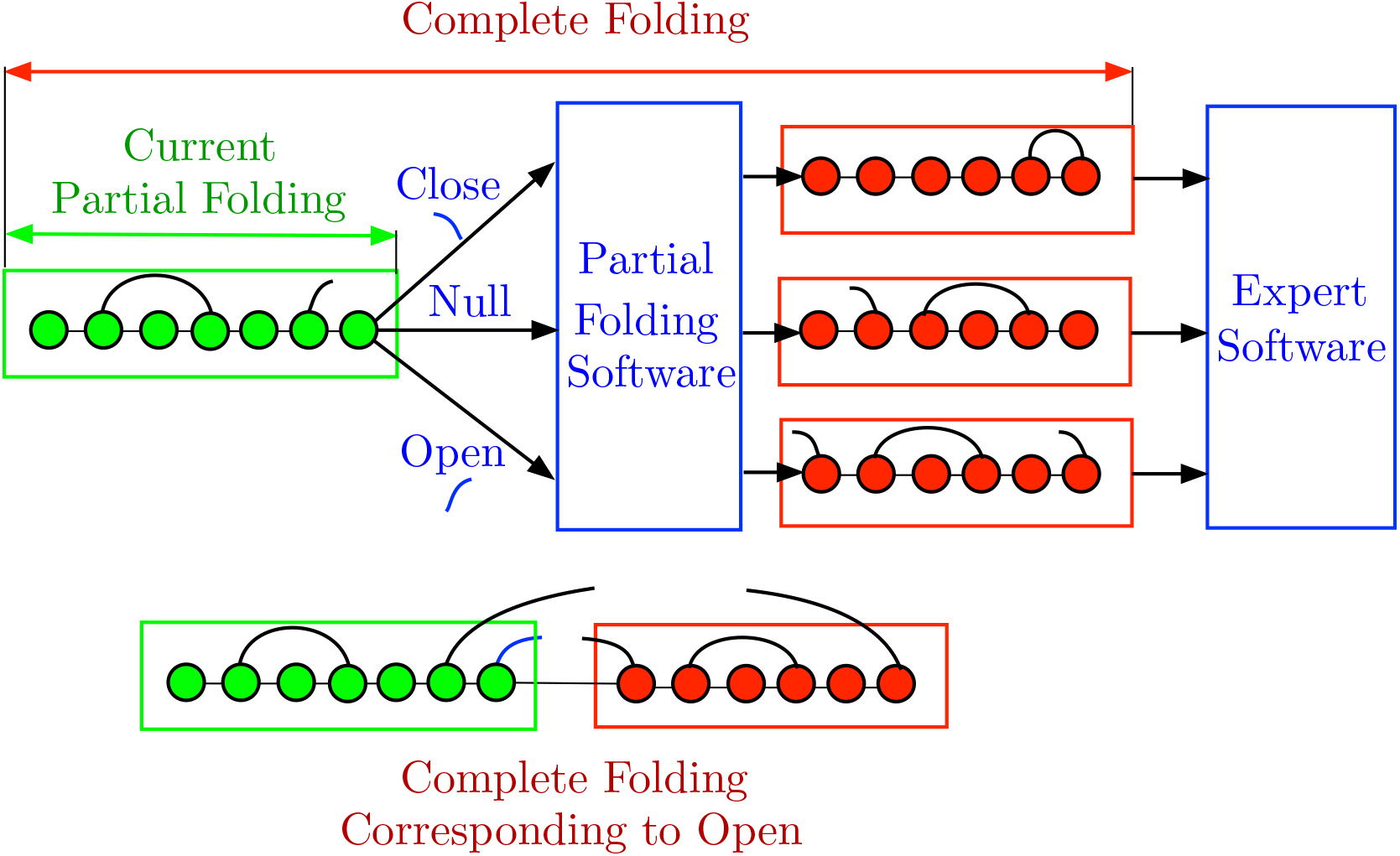
Overview of the proposed Expert-RNA.

#### 3.2.1 Rollout method

The rollout method was first introduced for the solution of discrete optimization problems in the paper by Bertsekas, Tsitsiklis, and Wu (see (Bertsekas et al., 1997)) and the book by Bertsekas and Tsitsiklis (see (Bertsekas and Tsitsiklis, 1996)). Rollout aims to find a solution of the general problem of minimizing a function *F*(*u*) over all *u* = (*u*_1_,…, *u^N^*) satisfying certain constraints. The components *u*_1_,…, *u_N_* can take a finite number of values, so this is a difficult discrete optimization. Rollout aims to find a suboptimal solution which improves over the solution produced by a given algorithm, called the base heuristic (in our case the base heuristic is RNAfold, but any other folding software can be used). This is accomplished by a sequence of minimizations of cost functions that are defined by *F*(*u*) and by partial soluiions produced by the base heuristic.

Our assumption is that, given any partial sequence 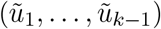, and any feasible value *u_k_*, the base heuristic produces a feasible complete solution of the form

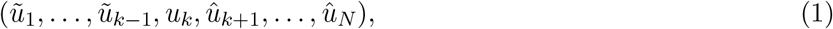

by constructing the complementary sequence (*û*_*k*+1_,…, *û_N_*) based on the knowledge of 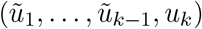. We also assume that the base heuristic can be applied to generate a feasible solution

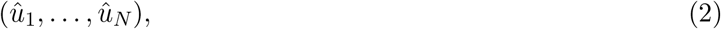

starting from the “empty” partial sequence. Initially, the rollout algorithm uses the base heuristic to generate, for each feasible value *u*_1_, the complementary sequence (*û*_2_,…, *û_N_*), and computes the initial solution component 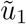 as

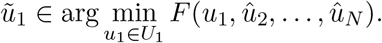

Where *U*_1_ is the set of feasible solutions at the first iteration. Then, sequentially for every iteration *k* ≥ 2, given the partial solution 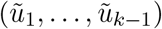, the rollout algorithm, considers all feasible values of *u_k_* and applies the base heuristic to generate the complete solution

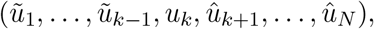

cf. Eq.(1). It then computes the value of *u_k_* that minimizes the cost function over all these complete solutions:

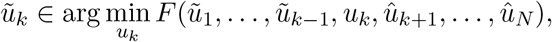

and fixes *u_k_* at the computed value 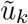. It then repeats with 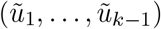 replaced by 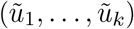. After *N* steps the rollout algorithm produces the complete sequence

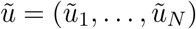

which is called the *rollout solution*. The fundamental result underlying the rollout algorithm is that under certain assumptions, we have *cost improvement*, i.e.,

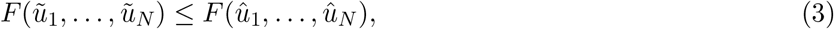

where (*û*_1_,…, *û_N_*) is the solution produced by the base heuristic starting with the empty partial solution (cf. Eq.(3)). Even when the assumptions needed for cost improvement are not satisfied, a simple modification, the so-called *fortified rollout algorithm*, produces a modified sequence that satisfies the cost improvement property in equation (3).

In addition to the fortified, several other versions of the rollout algorithm have been proposed in the literature; we refer to the reinforcement learning textbook by Bertsekas (see (Bertsekas, 2019)) and the monograph (see (Bertsekas, 2020)) for a detailed account, which includes discussions of rollout algorithms that incorporate constraints. Another version that is relevant to this work is *rollout with an expert*, which applies to problems where we do not know the cost function *F* of the problem, but instead we have access to an expert that can rank any two feasible solutions 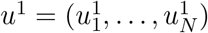 and 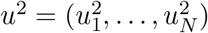 by comparing their values *F*(*u*^1^) and *F*(*u*^2^) (see (Bertsekas, 2019), Sect. 2.4.3, and (Bertsekas, 2020), Sect. 2.3.6). Still another version that is relevant to this work is rollout with multiple heuristics, which allows to use multiple base policies simultaneously, and also a variant that maintains multiple partial solutions simultaneously, and selectively enlarges some of these partial solutions.

#### 3.2.2 ExpertRNA Detailed Description

In this paper, we have used several algorithmic variants involving fortified rollout with multiple heuristics and multiple partial solutions. The expert that can rank two solutions is provided by the software package ENTRNA. The base heuristics are provided by the software package RNAfold (Sect. 2). It is important to note, however, that any expert software and base heuristic software are allowed within our algorithmic framework, subject to relatively weak restrictions (see the books (Bertsekas, 2019) and (Bertsekas, 2020)). In the current formulation of the rollout algorithm, we require the folder to be able to start from a partially formed structure and furthermore impose that during the folding, a selected base has to end up base-paired. Currently, only RNAfold allows for such a specific formulation of constraints. In the future version of our algorithm, we will modify the action definition so that it can work with folders that require to specify which base pairs are to be formed.

##### ExpertRNA main procedure

In the following, we provide the detailed iterations of ExpertRNA as a special implementation of a multi-branch fortified rollout algorithm.

###### Step 0. Initialization

provide the sequence of nucleotides **s** = (*s_k_*), *k* = 1, 2,…, *N* to be folded; initialize the current partial folding to be empty ***u***_0_ = (). Provide a folding software that, given a partial folding returns a complete structure. Provide an expert software with associated score function 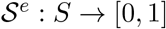, where *S* is the set of all feasible complete structures. Initialize the fortified solution 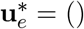. Set *k* ← 1;

###### Step 1. Move Generation

Given the partial solution (*u*_1_,…, *u_k_*), generate all feasible next assignments 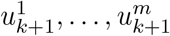, *m* ≤ 3 that satisfy the feasibility conditions (Sect. 3).

**For** each feasible move 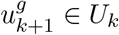, *g* = 1,…, *m*, where *U_k_* is the set of feasible actions at iteration *k* for the *k*^th^ nucleotide in the sequence:

###### Step 2. Structure Completion

Use the heuristic *H_k_* to generate the complete solution for the *g*^th^ feasible action:

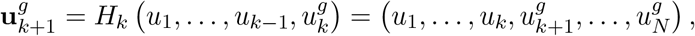

RNAfold (Sect. 2, (Lorenz et al., 2011)) in its constrained version, is used such that, considering the current state (i.e., the sequence of performed assignments up to the *k*^th^ iteration) and the next assignment 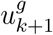 completes the structure based on minimium free energy (Zuker et al., 1999).

###### Step 3. Structure Evaluation

Score the candidate structures 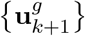, *g* = 1,… *m* using the expert scoring system 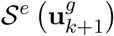 and save the top solution 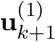. where 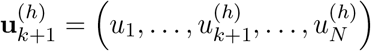, and the superscript (·) is to be interpreted as order index.

###### Step 4. Update the fortified solution

Update the fortified solution for the expert *e*: if 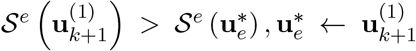, where 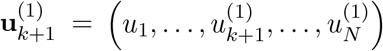.

*k* ← *k* + 1

###### Step 5. Stopping Condition

if *k* == *N*, STOP. Else Go to Step 1.

##### ExpertRNA extensions

A first extension to this basic algorithm we implemented is to maintain a set of *B* active branches. More specifically, the top-*B* branches are maintained at each point in time in the algorithm execution. Given that we have a maximum of *B_k_* · 3 feasible actions at each iteration *k*, where *B_k_* is the number of branches being maintained at iteration *k*, in general *B_k_* ≤ *B*. In particular, the set 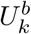 of feasible actions from a single branch *b* satisfies 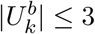. A second extension we implemented, with respect to the basic algorithm with multiple branches, was to allow for multiple experts. ExpertRNA is such that *B* is set to be a multiple of the number of experts. This choice is justified by the fact that no prior information is available that can help us choosing for which expert we should maintain more branches. Finally, in case multiple experts are adopted, a number of fortified solutions bounded by the number of experts is maintained at each iteration. The fortified solution(s) is updated at iteration k, if a structure with higher reward is identified for any of the experts. Then, at each iteration, a solution generated by a feasible action is compared to the fortified solution. If no action achieves a better score, the fortified action is chosen instead.

##### ExpertRNA move generation

A focal aspect of ExpertRNA is the generation of the set of feasible actions at iteration *k, U_k_* (Step 1, Move Generation in the general ExpertRNA algorithm). In the following, we detail the procedure for the generation of the feasible actions.

###### Step 0. Initialization

Iteration index *k*, partially folded structure *u*_1_,…, *u_k_*, and the unfolded nucleotide sequence 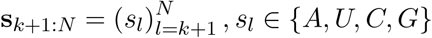. Initialize the set of feasible actions to the empty set 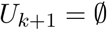.

###### Step 1. Action Generation

A feasible action represents the pair of nucleotide and the modality used to attach the nucleotide to the rest of the sequence. There are three possible actions: (i) open base pair, which we will refer to as 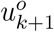; (i) close base pair, which we will refer to as 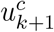; (ii) and the null action (i.e., simple sequencing), which we refer to as 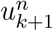.

###### Step 3. Feasibility Certification

We will verify whether the actions are feasible.

###### Open Base pairing

Condition 1: Given the incomplete structure **u**_*k*_, derive 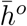, i.e., the number of open base pair that have not been closed. If the size of the remaining sequence satisfies 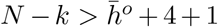, then condition 1 is satisfied, and we generate the set of candidate paired bases for the (*k* + 1)^st^ base;

Condition 2: If, among the nucleotides to pair, there exist at least one nucleotide that can physically pair with 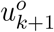, then condition 2 is satisfied;

Condition 3: If all the bases with open brackets 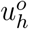, *h* = 1,…, *k* can close if the (*k* + 1)^st^ base is not a closed bracket, then condition 3 is satisfied;

If all conditions are satisfied 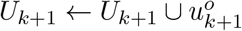.

###### Close Base pairing

Condition 1: Given the incomplete structure **u**_*k*+1_, derive 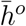, i.e., the number of open base pair that have not been closed, if 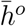. If 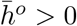, condition 1 is satisfied.

Condition 2: If there is an open base 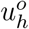, *h* ≤ *k* — 4 and the nucleotide is compliant with the current base, Condition 2 is satisfied.

If all conditions are satisfied 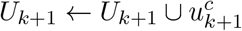.

###### Null pairing

Condition 1: Given the incomplete structure **U**_*k*_, derive 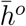, i.e., the number of open base pair that have not been closed, if 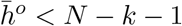, then condition 1 satisfied.

Condition 2: Calculate the number of nucleotides of the whole chain minus the position of last closed nucleotide, denote this number as **r_k_**. If 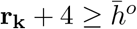, Condition 2 is satisfied.

Condition 3: If the (*k* + 1)^th^ nucleotide is unpaired, check whether the whole chain can be completed complying with feasibility constraints, which means all remaining incomplete open base pairs within the partial chain **u**_*k*_ can be paired using nucleotides which have not yet been assigned (A, U, C, G pairing constraints considered). If so, Condition 3 is satisfied.

If all conditions are satisfied 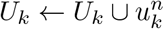.

**Figure 4:**
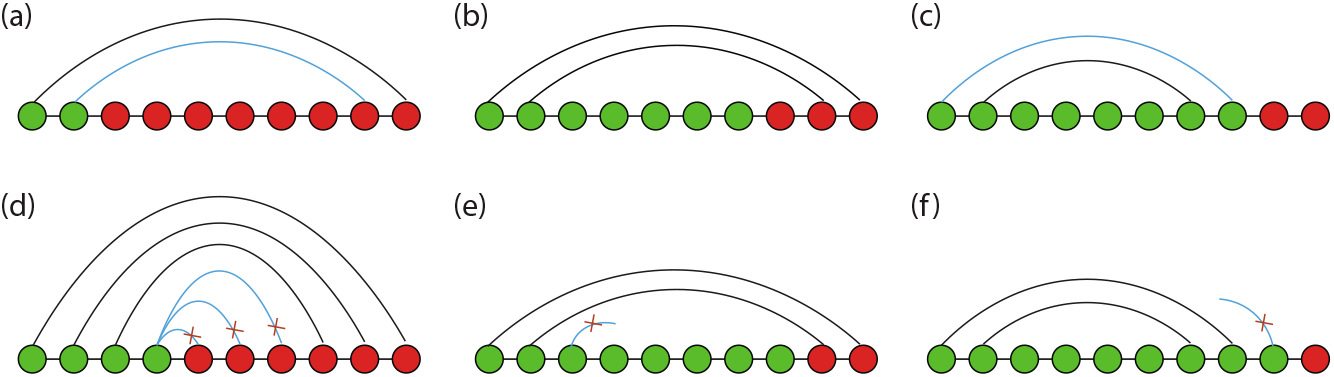
Examples of feasible and infeasible conditions for action as 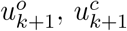, and 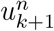. The green circles are the bases within the folded partial chain (the last green circle is the position we are currently assigning) and the red part is the unfolded part. The black arcs are implemented pairings whereas the blue arcs are being tested for feasibility. **(a)** the action 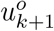 is feasible. **(b)** simple sequencing 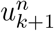 is feasible. **(c)** closing a base pair 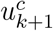 at the position is feasible. **(d)** opening a base pair 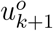 at the position is infeasible, since open base pairing condition 1 is violated. **(e)** sequencing 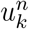 at the position is infeasible, since null pairing condition 1 is violated. **(f)** closing the base pair 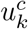 at the position is infeasible, since close base pairing condition 2 is violated.

## 4 Numerical Results

In this section, we compare the performance of ExpertRNA and RNAfold used in isolation. We highlight that judging the quality of a proposed structure is generally not a trivial problem, and this is the main motivation at the basis of data-driven approaches that allow to consider rewards different from free energy. In this analysis, we will consider the sequence-structure pairs in the data sets as the “ground truth”. Under this working assumption, the better method is the one which produces a sequence-structure pair that is more similar to the one in the data base. Similarity measures will be discussed.

### 4.1 Experimental Setting

#### Data set for training

When training ENTRNA, we adopted 1024 pseudoknot-free RNA molecules from the RNASTRAND data base (Andronescu et al., 2008). The length of the sequences within the data base ranges from 4 to 1192 nucleotides (Fig. 5(a) shows the distribution of the sequence length).

**Figure 5:**
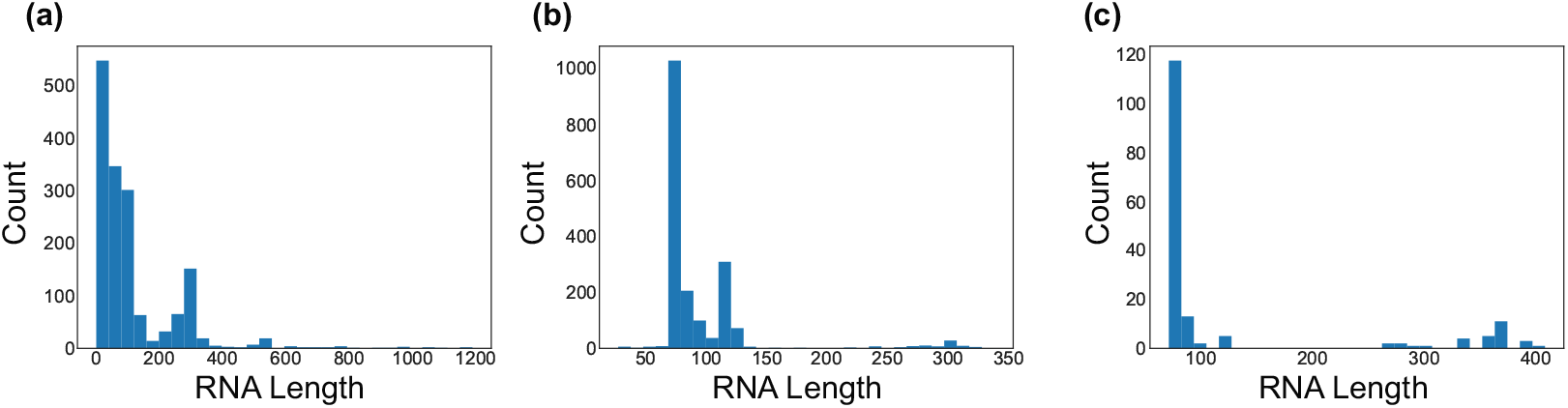
Distribution of RNA length in (a) RNASTRAND, (b) Mathews, and (c) Rfam.

#### Data set for testing

ExpertRNA was tested on two data sets. The first data set was obtained from the Rfam data base (Burge et al., 2013) by randomly selecting a subset of seed structures from distinct ncRNA families. Specifically, we used 167 sequences of length ranging from 71 to 408 nucleotides (Fig. 5(c)). The second data set is the benchmark data set of the RNAStructure tool (Reuter and Mathews, 2010), as populated by Mathews’ lab (Ward et al., 2017), which consists of natural RNA sequences with known secondary structures, comprising 1880 sequences of lengths ranging from 28 to 338 nucleotides (Fig. 5(b)). **Metrics** As previously mentioned, in this analysis, we consider better the algorithm that produces a structure that is close to the one in the data base. More specifically, to evaluate the quality of the produced structures, we look into three indicators:

- Free Energy (FE): Free Energy (FE) is a standard metric to evaluate the quality of an RNA structure. RNAfold is a FE minimizer. Hence, the ExpertRNA solution may have higher associated FE. The free energy calculation used here is from the ViennaRNA package and is the same one used by RNAfold (Lorenz et al., 2011);
- Matthews Correlation Coefficient (MCC): MCC (Matthews, 1975) is a popular metric in the RNA structure prediction field used to score the confusion matrix of a binary prediction. It takes into account the number of correctly and incorrectly predicted paired/unpaired bases and returns a score between −1 and 1, where −1 is a totally incorrect structure, 0 is the expected value of random assignment and 1 is a totally correct structure. The formal definition is:

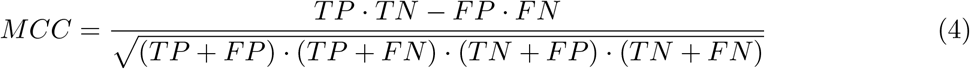

where TP, TN are the correctly identified paired and unpaired bases, respectively. FP, FN are the incorrectly paired and unpaired bases, respectively.
- Foldability: This metric is a score in the interval [0,1] quantified by ENTRNA. The higher the foldability the more likely is the sequence-structure pair to fold (Sect. 2).

#### ExpertRNA Settings

We created an extension of ENTRNA for this work, namely, the *No Free Energy ENTRNA* (*ENTRNA-NFE*). Specifically, we re-trained the ENTRNA classifier removing the free energy from the set of features used to train the Support Vector Machine at the basis of the expert. The rationale behind this modification was to allow to search for solutions (structures) that do not necessarily have low free energy. ENTRNA-NFE was a better-performing expert than the original ENTRNA as well as the ExpertRNA using *both* ENTRNA and ENTRNA-NFE simultaneously. We ran these several variants of ExpertRNA maintaining four branches at each iteration. In the case where we used both experts, we allowed each expert to maintain its two highest scoring branches at each iteration. As a result, at each iteration *k* of ExpertRNA, we have a number *B* = 4 of active branches and *B* = 4 structure foldings are performed. To reduce the computational demand, the foldings can be parallelized. All results reported were obtained from ExpertRNA with ENTRNA-NFE expert. Nonetheless, similar performance were obtained with the ENTRNA expert and when using both experts together.

### 4.2 Performance analysis

In the following, we analyse first the MCC as main discerning metric to individuate the characteristics of the proposed algorithm against the state of the art RNAfold. We then look into free energy and foldability to provide additional insights on the difference between the two approaches.

#### MCC Analysis

Tables 1–2 show the aggregate results in terms of the returned MCC. In particular, for each instance within the data base, we ranked the 4 resulting branches by foldability (best to worst) and report the average MCC, and median MCC 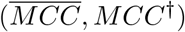 across the data set for the each branch. The average improvement in MCC over the RNAfold prediction 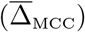, the standard deviation of the improvement (σ(Δ_MCC_)), and the pvalue associated to the hypothesis 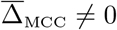, computed with a two-sided t-test. Here, Δ_MCC_ = MCC(ExpertRNA) – MCC(RNAfold) and we want this to be positive. Two main observations can be drawn from the analysis of the aggregate data. First off, the first branch solutions are always showing positive 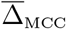, second, it is apparent that Expert RNA has a better performance over RNAfold in the Rfam data set as compared to the results from the Mathews data set.

**Table 1:** Aggregate MCC results produced by ExpertRNA for the Rfam data set.

| branch | $\overline{MCC}$ | $MCC^\dagger$ | $\overline{\Delta}_{MCC}$ | $\sigma(\Delta_{MCC})$ | p-value |
| --- | --- | --- | --- | --- | --- |
| 1 | 0.1017 | 0.1476 | 0.0558 | 0.1512 | 0.0492 |
| 2 | 0.1132 | 0.1597 | 0.0673 | 0.1516 | 0.0198 |
| 3 | 0.1001 | 0.1479 | 0.0542 | 0.1485 | 0.0553 |
| 4 | 0.1142 | 0.1588 | 0.0627 | 0.1500 | 0.0269 |

**Table 2:** Aggregate statistics on prediction quality by ExpertRNA for the Mathews data set.

| branch | $\overline{MCC}$ | $MCC^\dagger$ | $\overline{\Delta}_{MCC}$ | $\sigma(\Delta_{MCC})$ | p-value |
| --- | --- | --- | --- | --- | --- |
| 1 | 0.4327 | 0.5015 | 0.0066 | 0.1906 | 0.6378 |
| 2 | 0.4246 | 0.4984 | -0.0016 | 0.1927 | 0.9098 |
| 3 | 0.4211 | 0.4940 | -0.0051 | 0.1924 | 0.7123 |
| 4 | 0.4238 | 0.4966 | -0.0052 | 0.1943 | 0.7067 |

Fig. 6 shows the disaggregated MCC results. Two interesting observations emerge from the MCC distributions. First, for the Rfam data set (Fig. 6(a)-(b)), RNAfold never correctly identifies the correct structure, while ExpertRNA gets the complete or almost complete structure in a couple of cases. In Fig. 6(b), for the Rfam data set, the distribution of scores is shifted towards the right for ExpertRNA compared with RNAfold alone, with ExpertRNA performing worse in very few cases, indicating a better performance for ExpertRNA. The results are less clear-cut for the Mathews data set (Fig. 6(c) and (d)), where ExpertRNA does generally better on structures that RNAfold does very poorly on. However, there is also a significant population of structures that RNAfold gets correct or almost correct while ExpertRNA misses. These two features balance each other out resulting in no substantial improvement for ExpertRNA. In general, both ExpertRNA and RNAfold perform better on the Mathews data set compared with the Rfam data set.

**Figure 6:**
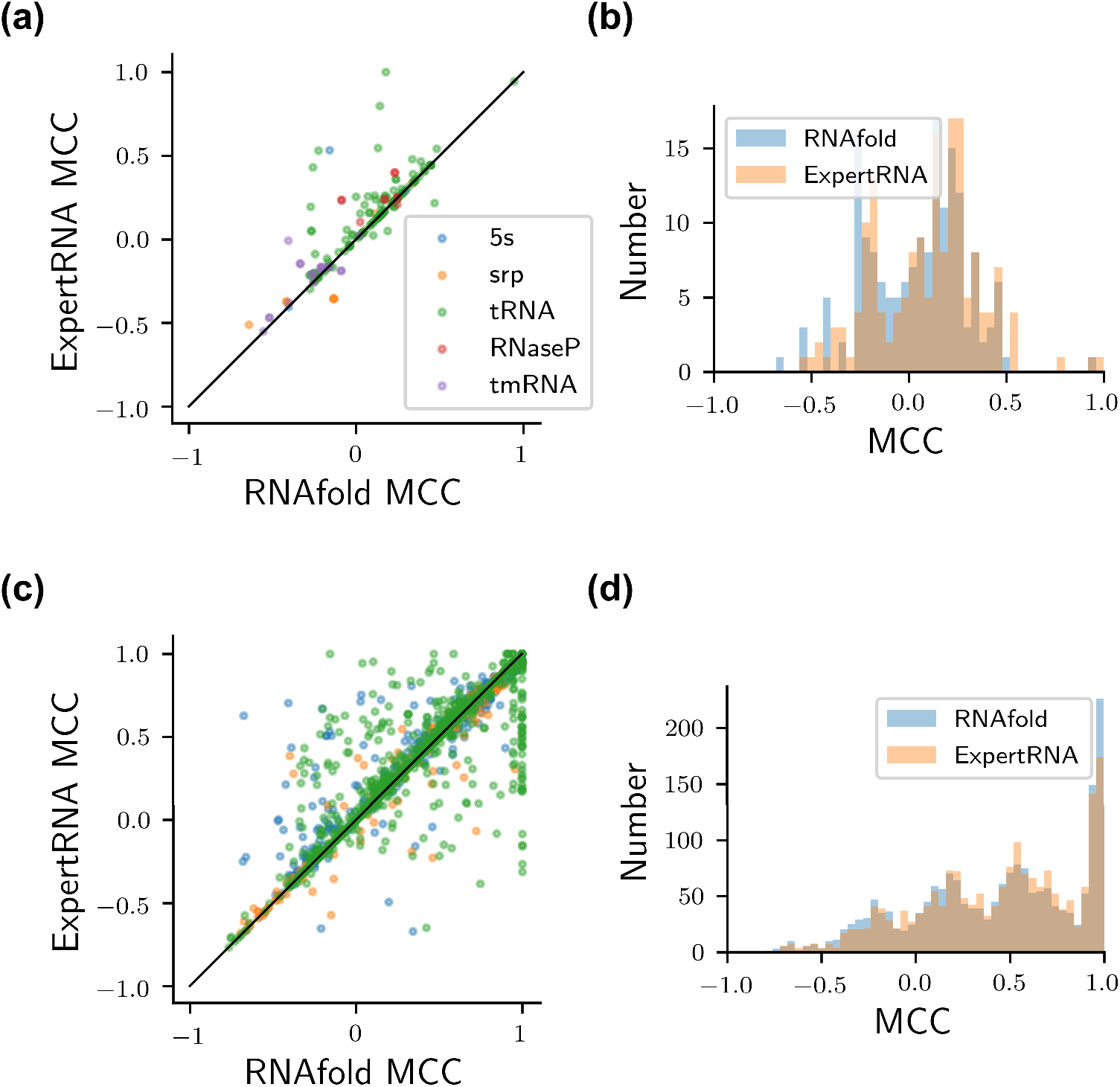
Performance of ExpertRNA compared with RNAfold on the two data sets broken down by RNA class. **(a)** and **(c)** Show the MCC scores for the structures generated from sequences in the Rfam database and Mathews database, respectively. **(b)** and **(d)** Show the distribution of MCC scores for the Rfam and Mathews database, respectively. In the scatter plots, the line of equal scores is also plotted (thin black line), as a reference.

#### Minimum Free Energy Analysis

Fig. 7(a)-(b) and Fig. 8(a)-(b) show how both ExpertRNA and RNAfold-generated structures generally have lower FE, as calculated by RNAfold, than the actual structure. Fig. 7(c) and Fig. 8(c) show that RNAfold tends to return lower FE than ExpertRNA. This is expected as RNAfold optimizes this value and it also reflects the result that ExpertRNA is able to predict non-MFE structures that are often more accurate by leveraging other measures of structure quality, such as the knowledge-based metrics considered by ENTRNA.

**Figure 7:**
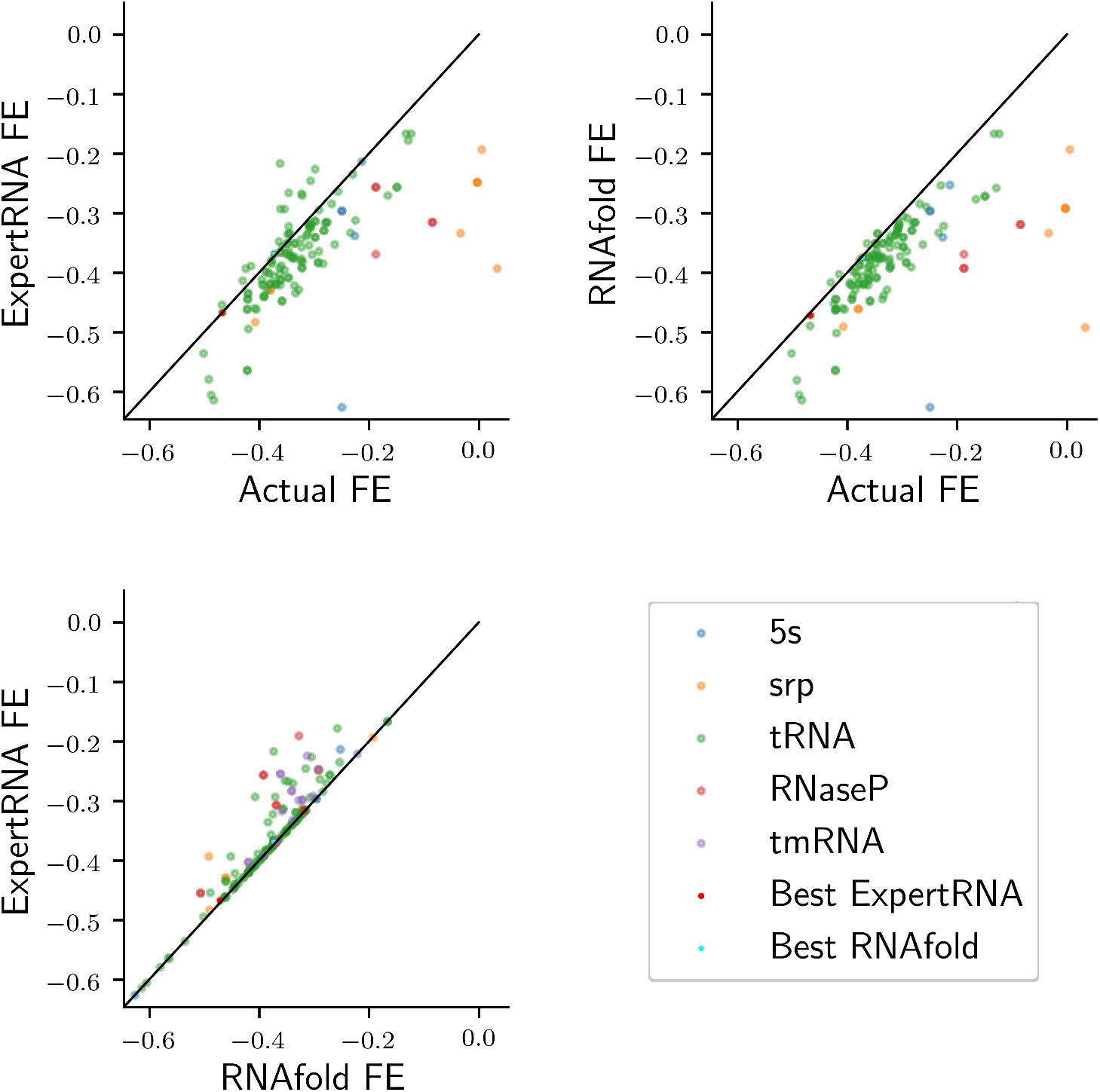
Free energy (FE) per nucleotide scores of structures from the Rfam data set as calculated by RNAfold for the actual structure, the predictions from ExpertRNA, and RNAfold. In each graph, input sequences are broken down by sequence type. Sequences where ExpertRNA did particularly well (MCC >= 0.9) are highlighted with red pips; there were no structures in this data set where RNAfold performed well. The line of equal scores is also plotted (thin black line) as a reference. **(a)** ExpertRNA-predicted structures compared with the actual structure. **(b)** RNAfold-predicted structures compared with the actual structure. **(c)** ExpertRNA-predicted structures compared with RNAfold-predicted structures.

**Figure 8:**
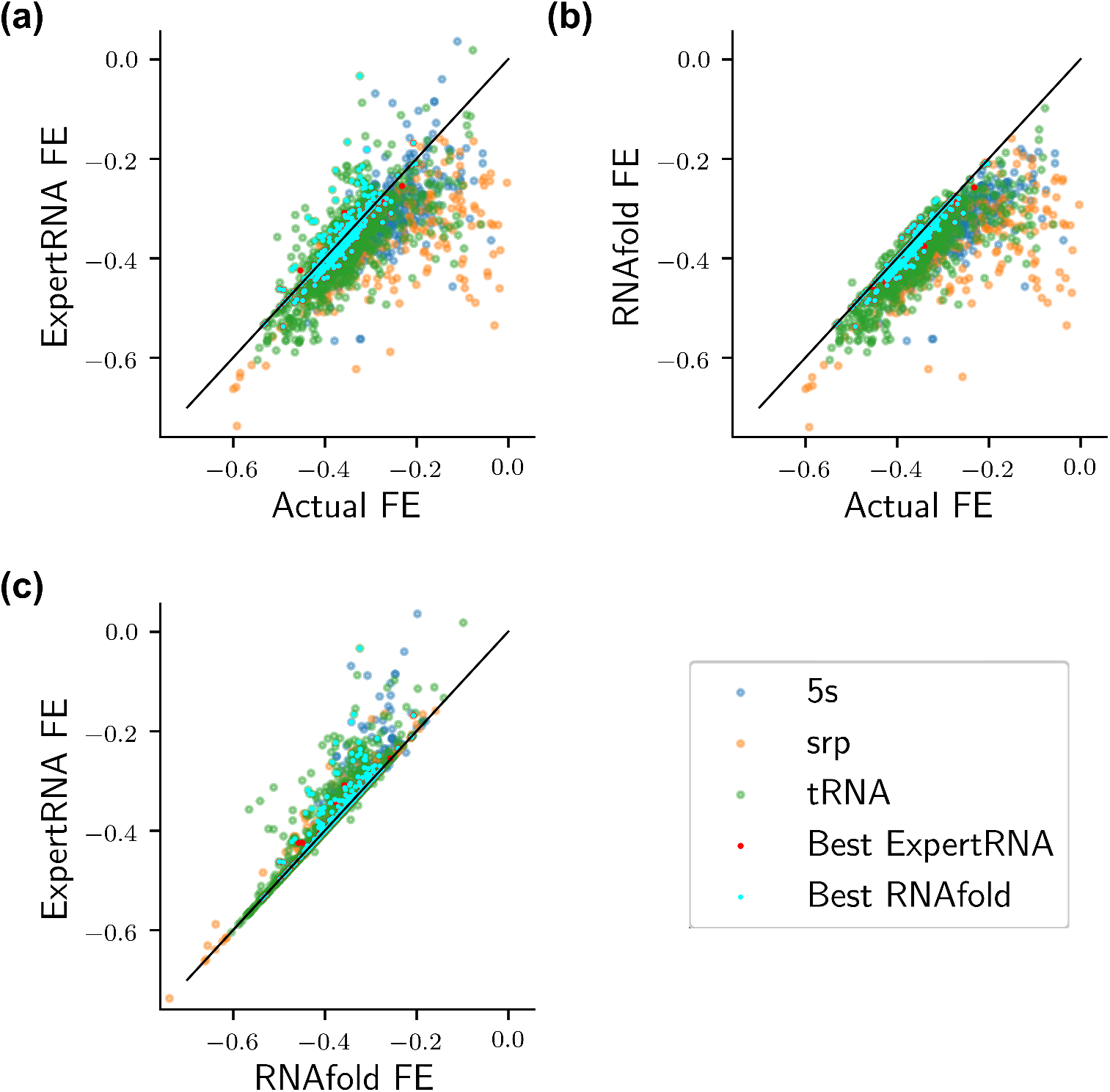
Free energy (FE) of structures from the Mathews database as calculated by RNAfold calculated for the actual structure, and the predictions from ExpertRNA, and RNAfold. In each graph, input sequences are broken down by sequence type. Sequences where ExpertRNA and RNAfold did particularly well (MCC >= 0.9) are highlighted with smaller cyan and red pips, respectively. The line of equal scores is also plotted (thin black line) as a reference. **(a)** ExpertRNA-predicted structures compared with the actual structure. **(b)** RNAfold-predicted structures compared with the actual structure. **(c)** ExpertRNA-predicted structures compared with RNAfold-predicted structures.

#### Impact of the Expert Score

An important element of ExpertRNA is the expert adopted. While, in principle, the algorithm can make use of any expert, it is important to understand the impact of the adopted expert on the solution. As detailed in Sect. 2, in this implementation, the adopted expert is a machine learning algorithm that, given a set of sequence-structure pairs returns a score for each of them based on how “likely” the pairs are, where such likelihood is referred to as *foldability* in the original paper (Su et al., 2019). As shown in Fig. 9 and Fig. 10, as expected, ExpertRNA predictions have the higher foldability. The fact that ExpertRNA performed better on the Rfam data set suggests that ENTRNA foldability is a slightly better expert than RNAfold’s MFE measure. However, the many incorrect structures with higher foldability than the actual structures points to the need for more research in developing better performing expert software.

**Figure 9:**
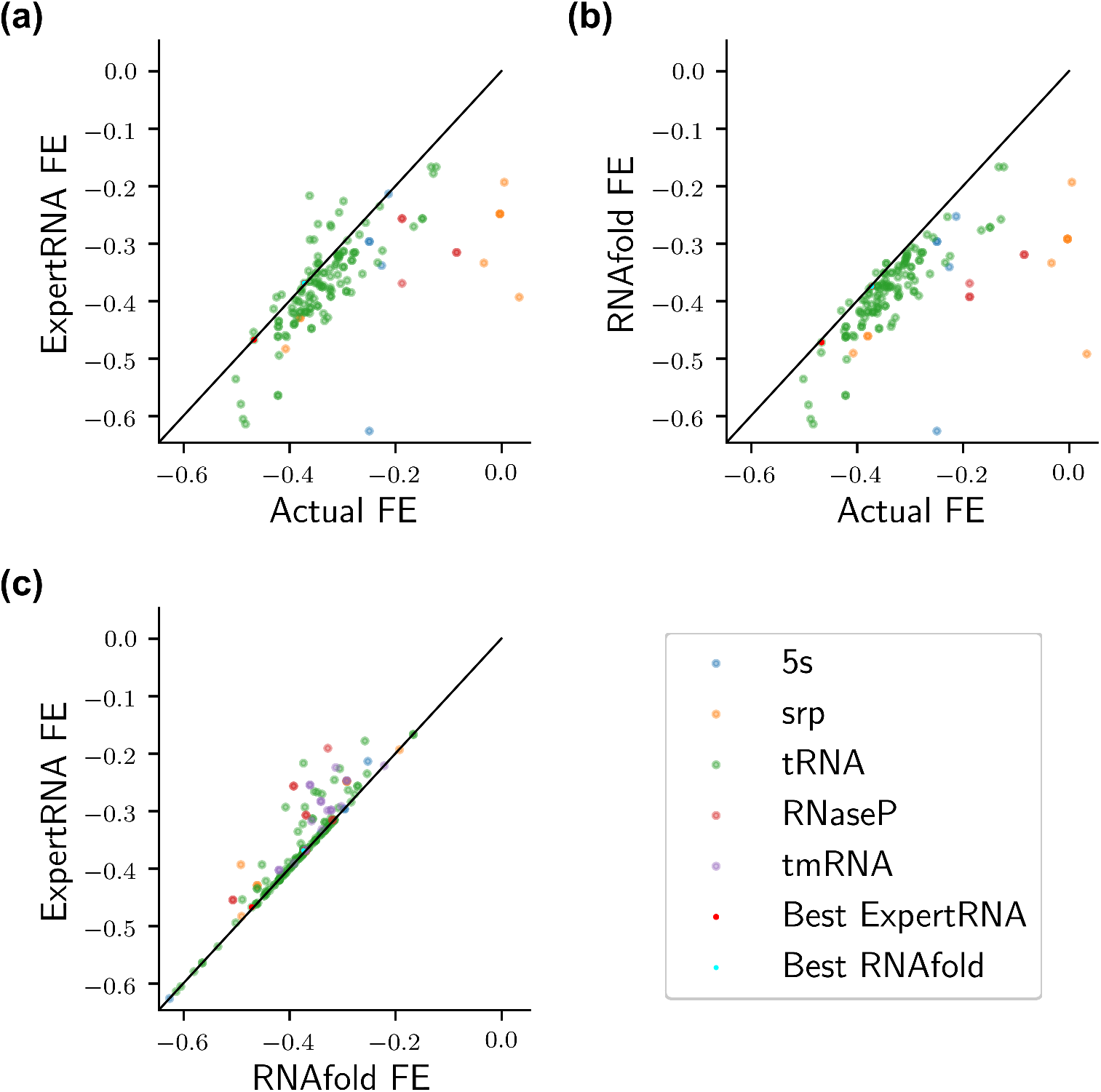
Foldability scores for structure predictions from the Rfam data set as calculated by the expert, ENTRNA-NFE. In each graph, input sequences are broken down by sequence type. Sequences where ExpertRNA did particularly well (MCC >= 0.9) are highlighted with red pips; there were no structures in this data set where RNAfold performed well. The line of equal scores is also plotted (thin black line) as a reference. **(a)** ExpertRNA-predicted structures compared with the actual structure. **(b)** RNAfold-predicted structures compared with the actual structure. **(c)** ExpertRNA-predicted structures compared with RNAfold-predicted structures.

**Figure 10:**
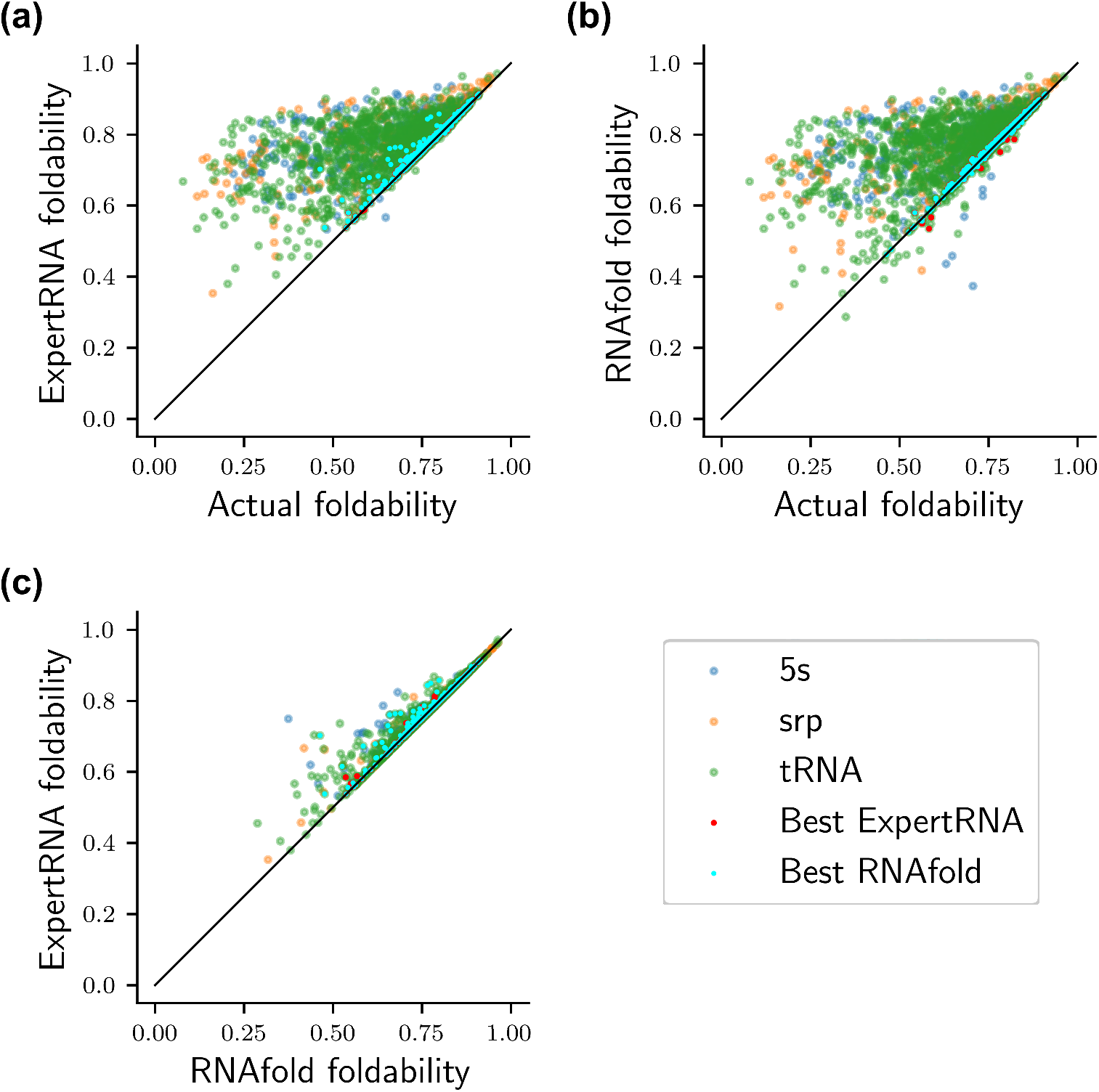
Foldability scores for structure predictions from the Mathews data set as calculated by the expert, ENTRNA-NFE. In each graph, input sequences are broken down by sequence type. Sequences where ExpertRNA did particularly well (MCC >= 0.9) are highlighted with smaller cyan and red pips, respectively. The line of equal scores is also plotted (thin black line) as a reference. **(a)** ExpertRNA-predicted structures compared with the actual structure. **(b)** RNAfold-predicted structures compared with the actual structure. **(c)** ExpertRNA-predicted structures compared with RNAfold-predicted structures.

#### Predicted Structures

We show a sample of the structures generated by ExpertRNA and compare them to the prediction of the RNAfold algorithm alone as well as the actual structure. In particular, we look into cases where the predicted structure generated by RNAfold significantly differs from the known structure. Visualizations were generated using the publicly available Forna software (Kerpedjiev et al., 2015). Fig. 11 shows the structures where RNAfold is particularly underperforming in the Rfam data base.

**Figure 11:**
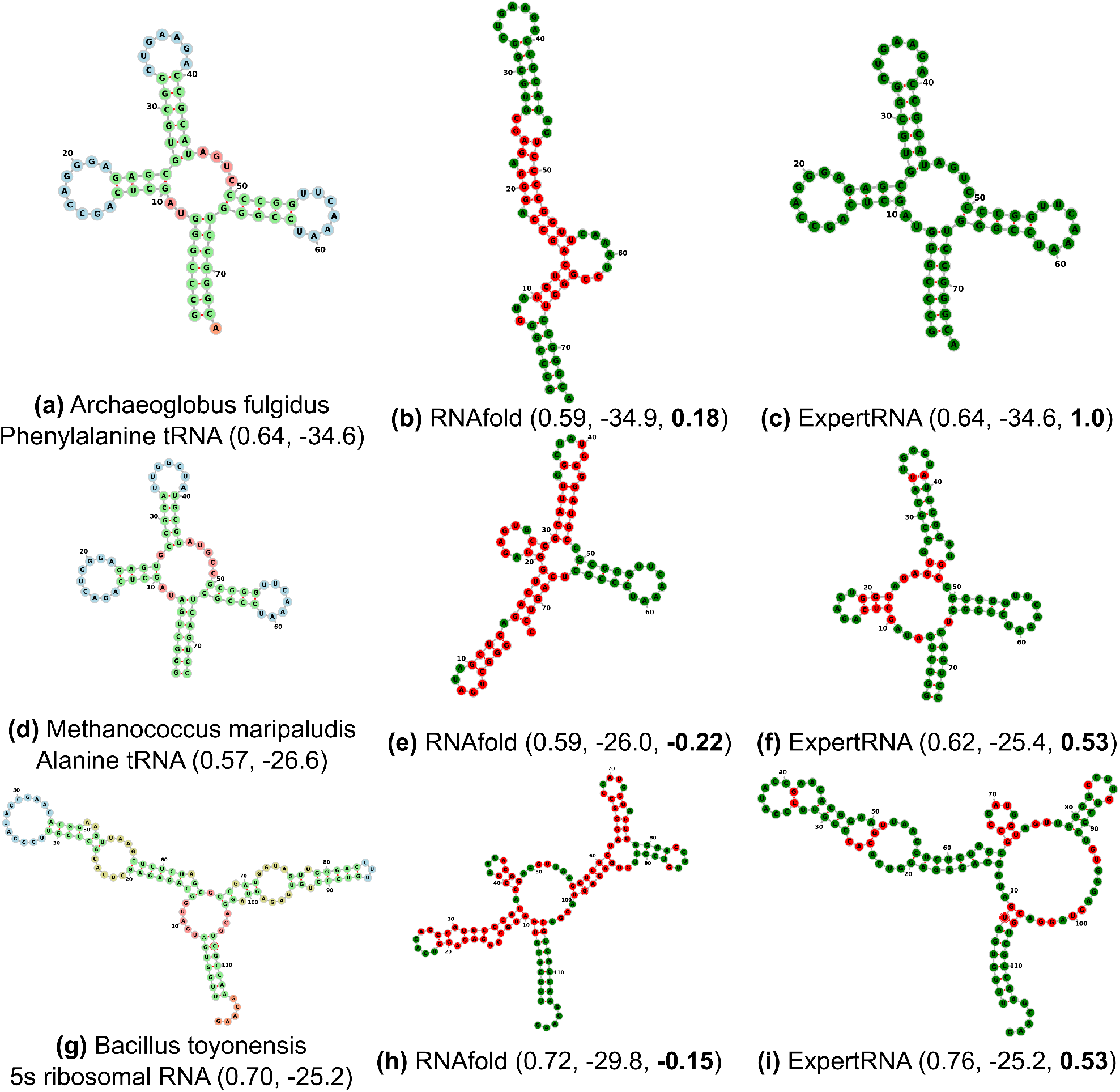
Example structure from the Rfam data set with the best performance improvement between ExpertRNA and RNAfold. Each row contains the actual structure (a-d-g), the RNAfold (b-e-h), and the ExpertRNA prediction (c-f-i). Nucleotides whose pairing was predicted incorrectly are shown in red in panels (b) and (c), while correct pairings are green. The numbers in parenthesis are: foldability of the structure as evaluated by ENTRNA-NFE, free energy as evaluated by RNAfold, and MCC compared with the actual structure (bold): **(a-c)** A phenylalanine tRNA from Archaeoglobus fulgidis; **(d-f)** An alanine tRNA from Methanococcus maripaludis; **(g-i)** A 5s rRNA from Bacillus toyonensis.

Fig. 12 shows the structures where RNAfold is particularly underperforming in the Mathews data set.

**Figure 12:**
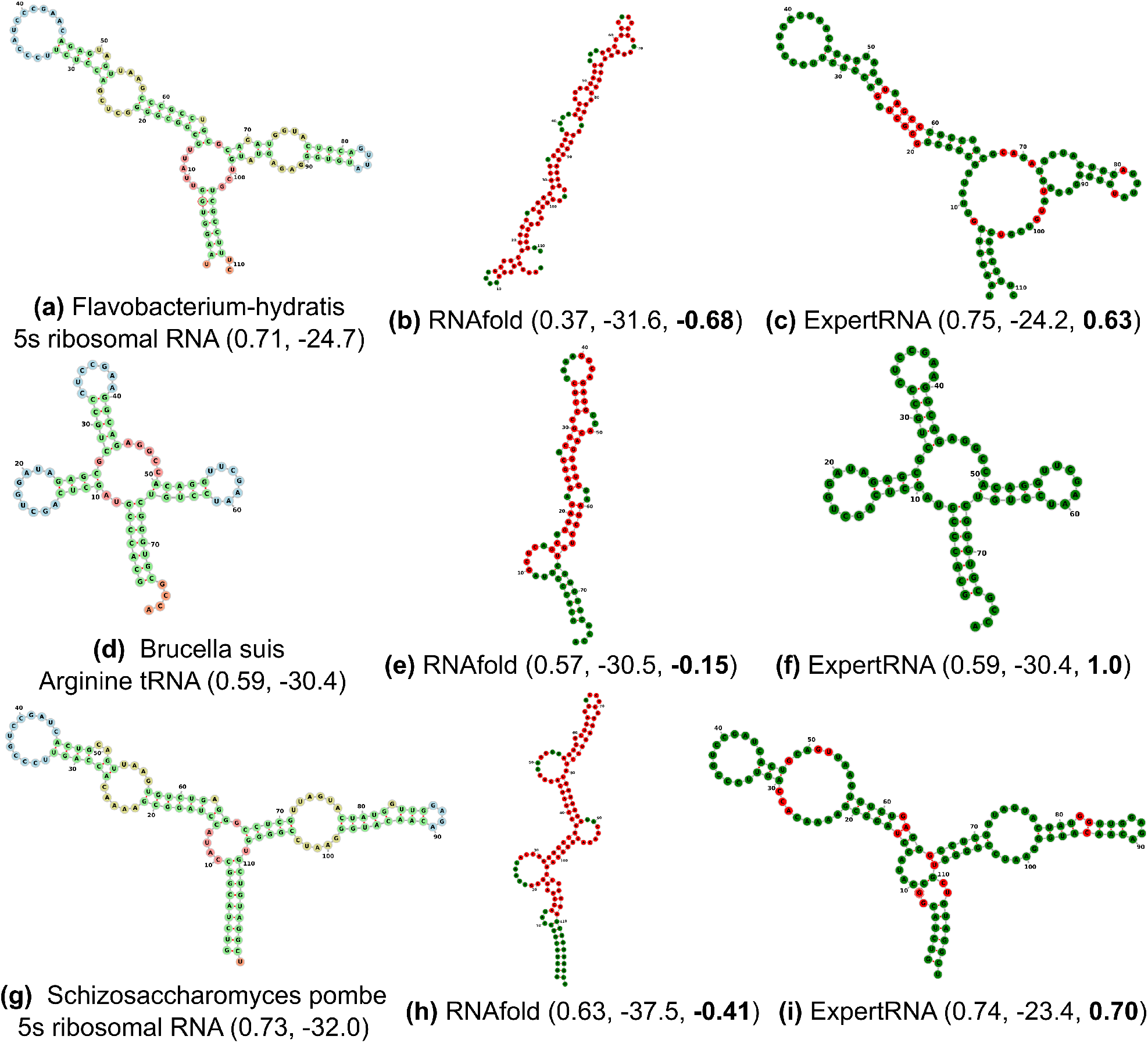
Example structure from the Mathews data set with the best performance improvement between ExpertRNA and RNAfold. Each row contains the actual structure (a-d-g), the RNAfold (b-e-h), and the ExpertRNA prediction (c-f-i). Nucleotides whose pairing was predicted incorrectly are shown in red in panels (b) and (c), while correct pairings are green. The numbers in parenthesis are: foldability of the structure as evaluated by ENTRNA-NFE, free energy as evaluated by RNAfold, and MCC compared with the actual structure (bold) **(a-c)** A 5s rRNA from Flavobacterium hydratis. **(d-f)** An agrinine tRNA from Brucella suis. **(g-i)** A 5s rRNA from Schizosaccharomyces pombe.

### 4.3 Computational Complexity

Another important aspect of structure folding is associated to the complexity of the algorithm. For the current implementation, ExpertRNA has a complexity 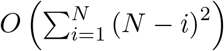, where (*N* – *i*)^2^ is the RNAfold complexity for a sequence of size (*N* – *i*). Such quadratic behavior is evident in Fig. 13 (the results are for the Mathews data set, and similar results were obtained for the Rfam data set). While computational efficiency is a known challenge in secondary structure prediction, a promising avenue will be to investigate multi-agent rollout (Bertsekas, 2020) as a way to boost the efficiency.

**Figure 13:**
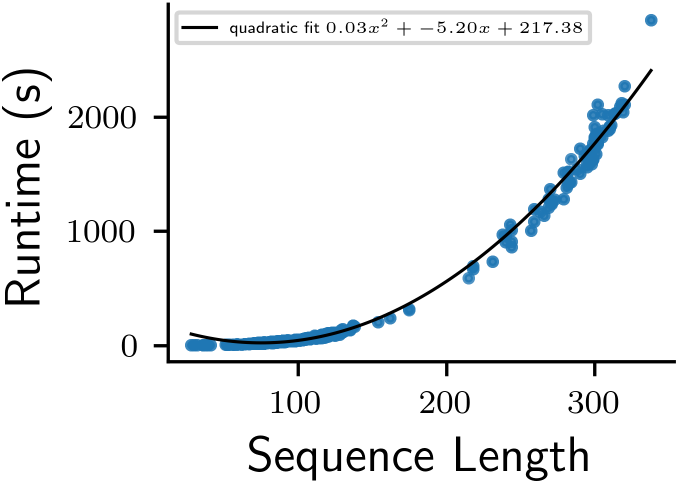
Computational effort for ExpertRNA across the Mathews data set with quadratic fit shown.

### 4.4 Summary

Compared to the RNAfold tool, ExpertRNA performs better on the data set of known structures from Rfam, and approximately equivalently on the data set from the Mathews lab. In several cases, the ExpertRNA structures had higher free energy (compared to RNAfold) according to the nearest-neighbor model. However, they are closer to the actual known structure. Structures predicted by both ExpertRNA and RNAfold almost always have lower free energy than the actual structure, further indicating the limitations of free energy approximations in identifying correctly folded structures. In most cases, where ExpertRNA found structure in better agreement with the actual known structure, the FE of that structure was only slightly higher than the FE of RNAfold. However, in few cases (e.g. Fig. 12(a)-(c),(g)-(i)), the solutions where ExpertRNA had much better agreement with the correct structure, RNAfold solution had much lower free energy. ExpertRNA hence shows the ability to identify the correct structure even if its predicted FE is quite higher than the MFE prediction by integrating other experts that may implicitly or explicitly account tertiary contact and kinetic information.

## 5 Conclusions

In this paper, we propose a new framework, ExpertRNA, for the automatic folding of secondary structures for RNA molecular compounds. ExpertRNA builds upon the fortified rollout algorithm and generalizes the architecture to allow for the consideration of multiple experts that can evaluate, at each iteration, the solutions generated by the base heuristic. ExpertRNA allows to control the growth of the rollout branches by enabling only a fixed number of alternatives at each iteration to be maintained. This differs from the traditional parallel expert implementation where each expert is responsible for independent branches leading to the exponential growth of the maintained branches. Differently, ExpertRNA allows the folding algorithms to interact and the experts to evaluate all the solutions generated by all the folders at each iteration. This feature of ExpertRNA balances out different strengths and weaknesses of folding tools and experts. Numerical results show the advantage of the proposed approach especially over the data set where the heuristic alone does not perform well. We showed that, for several natural sequences, ExpertRNA was able to correctly predict secondary structures whose free energy was much higher than the MFE structure predicted by RNAfold.

Further investigation is being carried out to employ multiple different folding algorithms. We argue that, in the process of generating the solution, perhaps one of the folders will be performing very well, but, after some iterations, a different folder may produce superior results. In some sense, ExpertRNA tracks the best folder, online, as the sequence is being constructed. Also, we are investigating the opportunity to use more/different experts. In light of this, a possible future direction is to include more features in ENTRNA training or combination with additional expert scoring (e.g., based on covariation information of homologous sequences and SHAPE experimental data) would improve the prediction accuracy.

Finally, the introduced framework is not limited to RNA secondary structure and can be extended to *de novo* predictions of RNA tertiary structure or protein structures. Most of the currently used models for structure prediction allow to specify contact constraints that denote pairs of residues that are in contact. They are hence amenable to be used in ExpertRNA, which requires each folding tool to start from a partially formed structure. There are currently over 200 different methods that participate in the CASP challenge that compares different folding algorithms of proteins structure prediction (Moult et al., 2018), and over 10 different methods in its RNA counterpart, RNA-puzzle (Cruz et al., 2012), and these algorithms can be used as folders in the Expert-RNA framework. The experts in such implementation can correspond to the system energy calculated, e.g., with Rosetta (Rohl et al., 2004), and other knowledge-based potentials. The Expert-RNA tool is freely available at https://github.com/MenghanLiu212/RL-RNA, and the scripts used to process the output of Expert-RNA into Fig. 6–12 are freely available at https://github.com/ErikPoppleton/rollout_plotter.

